# Eduomics: a Nextflow pipeline to simulate -omics data for education

**DOI:** 10.1101/2025.11.24.690117

**Authors:** Lorenzo Sola, Davide Bagordo, Simone Carpanzano, Mariangela Santorsola, Francesco Lescai

## Abstract

Moving past learning just algorithms and code is a key challenge of bioinformatics education: the ideal goal is for students to acquire higher-order knowledge such as the ability to solve biological problems with the appropriate tools, and more importantly learn to interpret the results in the broader context where bioinformatics is needed. To design such a teaching and learning experience, data simulations play a key role: however, there is a massive barrier to adoption. Different data types are produced by different tools, requiring educators to learn each of them and adapt their workflow to the necessary dependencies, requirements and input files. Additionally, most existing data simulation solutions are meant for benchmarking and methods development rather than education and cannot provide the context needed to teach students the critical interpretation skills they need to move beyond problem-based learning to what we call storyline-based learning. A significant effort must be placed also when many datasets with the same characteristics are needed, such as in tutoring or assessment in higher education. Here, we present eduomics: a Nextflow pipeline meant to automate the simulation workflow for both genomic and transcriptomic next-generation sequencing data, and to produce realistic clinical scenarios to provide students with clues and a biomedical story necessary for the interpretation of their results. Eduomics removes barriers to adoption, by requiring the user to just decide which chromosome datasets should be simulated on, and which type of data they would like to simulate. There is no need to learn specific tools and resolve their dependencies. The use of Gemini API provides an innovative approach to generate plausible clinical scenarios, consistent with the genes where either a pathological mutation or differential expression has been simulated. With eduomics, we offer an accessible and scalable solution to design comprehensive learning experiences and innovate bioinformatics education.

**Author Summary:** Modern bioinformatics education faces a dual challenge: students must learn how to analyse data and interpret results within the biological problems they aim to solve, while instructors who wish to design such a comprehensive learning experience should introduce appropriately simulated data in their teaching. However, the complexity of existing simulation tools is an often daunting barrier to overcome: current solutions require different software for different data types and provide no support for teaching interpretation skills. Most available simulators were designed for benchmarking and method development, not for education: therefore they lack the biological and clinical context needed to move beyond the traditional problem-based learning.

To address these challenges, we developed eduomics, a Nextflow-based end-to-end pipeline that automates the simulation of genomic and transcriptomic data in a fully automated manner for educational use. Educators only need to choose the type of data to simulate, while the pipeline handles all underlying tools and dependencies. As a groundbreaking element, we integrated the Google Gemini API to generate patient-inspired clinical scenarios that gives students realistic clues and a biomedical narrative in which to interpret the results. This approach complements the traditional problem-based learning by offering a more complete, storyline-based learning experience. By combining automated data simulation and clinical storytelling, eduomics provides an accessible and scalable solution that support an engaging and immersive learning experience for both educators and students.

## Introduction

The rapid evolution of Next generation sequencing (NGS) technologies has profoundly reshaped both biomedical research and clinical practice. The continued decrease in sequencing costs, together with the proliferation of multi-omics data, is transforming the life sciences into a data-intensive discipline. As a results, students enrolling in biomedical studies are increasingly expected to manage complex computational workflows: bioinformatics has thus become a cornerstone of modern life sciences education. Consequently, there is a growing consensus that bioinformatics training should be introduced early in higher education curricula [1–3] to equip future scientists with the computational and analytical competencies required in contemporary laboratories.

In this context, simulated datasets offer a controlled and flexible environment where learners can engage with realistic workflows without raising ethical or privacy concerns. However, using existing tools for data simulation in an educational setting remains challenging. Most of these tools are designed for benchmarking software or algorithm development, rather than for educational purposes. An educator faces a steep learning curve which is a barrier to the adoption of simulations as part of common teaching and learning designs. Their use typically requires choosing the appropriate software, installing their dependencies, generating compatible reference datasets, creating input files in specific formats, and manually integrating intermediate outputs. Even when these obstacles are faced with, significant extra work is required to connect the data generated by existing simulators with the underlying molecular mechanisms and the clinical phenomena they participate in. However, embedding the simulations in the appropriate context is necessary to interpret the analysis results. This is an essential skill that educators aim to develop in students. A few examples of this gap can be observed in genomics, where different solutions have been developed to simulate genetic and transcriptomic data. For example, a broad ecosystem of variant-injection and genome perturbation tools enables users to embed predefined variants into simulated reads or alignment files, thereby providing high-fidelity benchmarks for variant calling and alignment evaluation [4–8]. Similarly, a variety of tools have been developed to simulate RNA sequencing (RNA-seq) reads for benchmarking quantification and differential expression workflows. This software can model experimental biases such as GC content, transcript length, and positional effects, enabling the creation of datasets with known differentially expressed genes (DEGs) [9–11].While these tools produce realistic data, their workflow is fragmented and context-dependent: each focuses on a single step of the process, leaving users responsible for chaining outputs, normalising file formats, and managing parameters. An additional challenge educators face, especially when dealing with data simulations in official curricula, is the need to generate hundreds of datasets with the same characteristics: this is because different datasets are needed not only for classes, but also for exercises, tutoring and more importantly assessments.

Automated, accessible, scalable and portable solutions are needed to overcome this key challenge. In recent years, a few end-to-end next-generation sequencing (NGS) simulation pipelines have emerged, which generate synthetic biological datasets designed to mimic sequencing experiments. These frameworks reproduce empirical features such as sequencing errors, mutation spectra, and gene expression profiles, producing realistic resources for benchmarking, workflow optimisation, and software validation [12–14]. Yet even these comprehensive pipelines remain primarily research- and benchmark-oriented: they offer technical realism without addressing the need, compelling in education, to contextualise the molecular data simulated within biological systems or even clinical scenarios.

To bridge these key gaps, we developed eduomics: the pipeline is designed to generate hundreds of datasets enabling the acquisition of both technical proficiency as well as conceptual understanding, and to provide teachers with a single end-to-end tool which transforms complex simulations into an accessible educational opportunity. Eduomics is a Nextflow-based pipeline specifically designed for educational use that automates all key steps of NGS simulation, from reference preparation to results interpretation. Unlike traditional simulators, this pipeline generates biologically coherent variant calling and RNA sequencing datasets enriched with clinically validated variants and realistic transcriptomic profiles. Each simulation is defined by a known ground-truth, ensuring full control over the injected variants or expression changes and allowing students to trace analytical outcomes back to their biological causes. By integrating Google Gemini’s generative capabilities, the pipeline links molecular alterations to AI-derived clinical narratives creating immersive learning experiences. This educational design goes beyond the traditional problem-based learning (PBL) by introducing a more complete, scenario driven framework. Through the combination of validated data, clinical context, and narrative design, this approach establishes a pedagogical framework grounded in storyline-based learning, enabling students to explore the biological significance of computational results in realistic biomedical contexts.

## Results

The eduomics pipeline has been developed using the Nextflow DSL2 language, combining modular processes in line with nf-core community standards. The pipeline integrates 16 modules from the shared nf-core repository and 14 modules specifically developed to enable the pipeline functionalities (called “local”, according to the nf-core community nomenclature).

The release 1.1.0 (https://zenodo.org/records/17552833 accessed on 7 November 2025) accepts a sample sheet in CSV format as an input: this file defines the parameters for each simulation, which produces validated variant calling and RNA-seq datasets together with AI-generated clinical scenarios. An overview of the main workflow is shown in Fig 1, and a detailed pipeline diagram is provided in S1 Fig.

**Fig 1.**
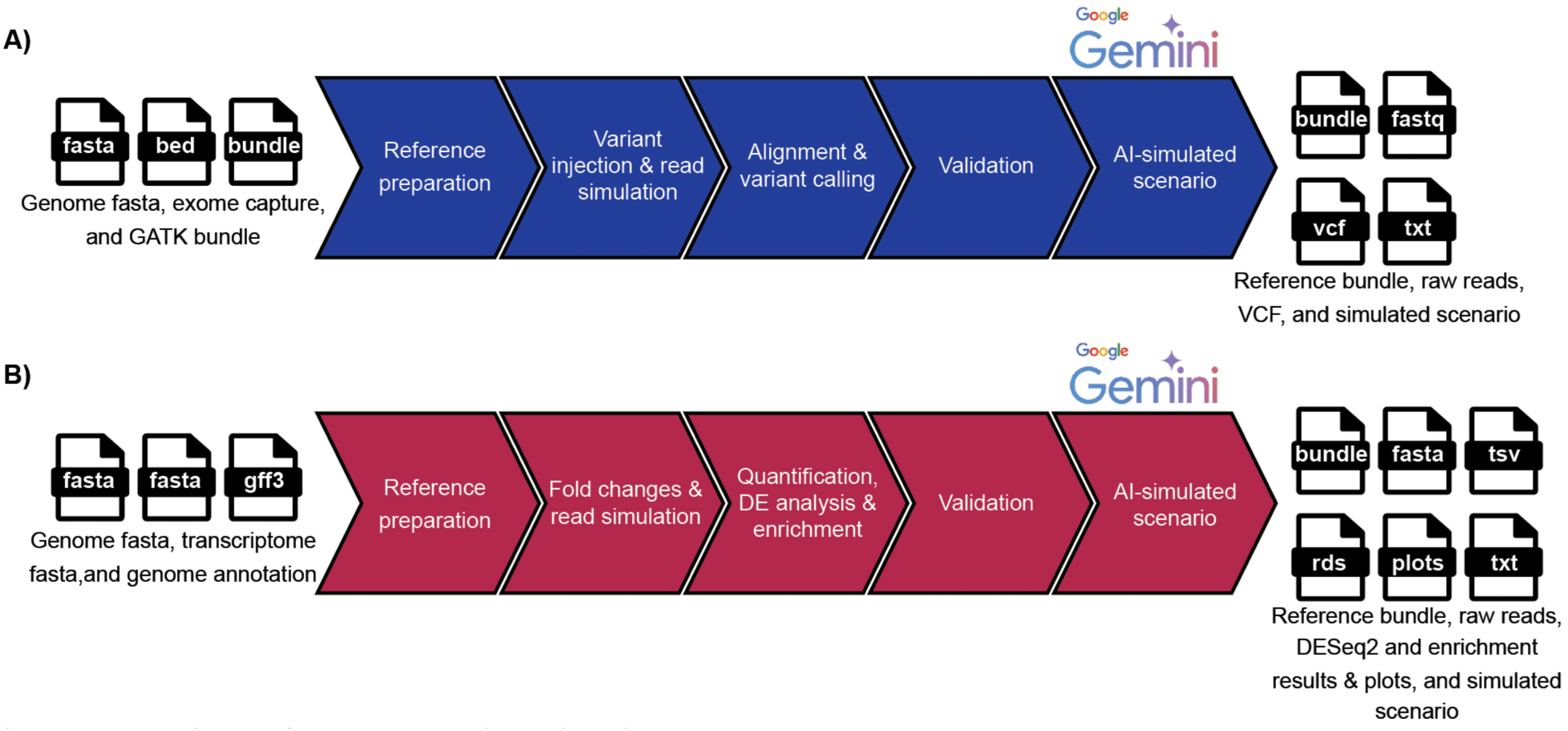
Overview of the eduomics pipeline. The pipeline comprises two main workflows: (A) the variant calling workflow, which simulates resequencing data with selected likely pathogenic and pathogenic variants, and (B) the RNA-seq workflow, which generates transcriptomic datasets with biologically meaningful differential expression. The outputs produced from each workflow converge into a final common module that employs Google Gemini to generate patient-inspired clinical scenarios. The arrows illustrate the logical progression of each workflow, from initial inputs to final outputs.

The variant calling workflow (Fig 1A) generates synthetic resequencing data by injecting known likely pathogenic or pathogenic variants from the ClinVar release into a user-defined genomic interval. This workflow generates simulated raw reads from a case/control sample design and processes them according to GATK Best Practices to validate the results. This ensures that the DNA datasets resemble realistic resequencing experiments while embedding variants of clinical relevance. The RNA-seq workflow (Fig 1B) simulates transcriptomic data with biologically meaningful differential expression. This workflow prepares the reference files, applies fold changes to selected gene sets, and generates synthetic read counts, followed by quantification and enrichment analysis, to validate the simulated data. This design allows the simulation of RNA-seq datasets that reproduce expression patterns consistent with biologically relevant pathways.

By processing both type of simulations through a complete analysis workflow, only those simulated datasets which produce the expected results are saved by the pipeline. Validated results serve as input to a common module that employs the Google Gemini API to generate patient-inspired clinical scenarios. These scenarios represent a key outcome of eduomics, with the goal of bridging the operational understanding of computational tools with the clinical interpretation as a key element of the teaching and learning approach.

To evaluate performance, we benchmarked eduomics on human chromosome 22. The pipeline was tested both on an on-premise HPC system (SLURM scheduler) and on Google Cloud, to demonstrate full portability across computational environments. As a case-study, we present here the run on Google Cloud Platform, which allows a more fine grained report of the workflow performance and resource utilisation (Fig 2), as well as the actual costs. The simulation completed in approximately seven hours (Fig 2A), involving a total of 11,794 submitted jobs (Fig 2B) and an estimated execution cost of €199,11.

**Fig 2.**
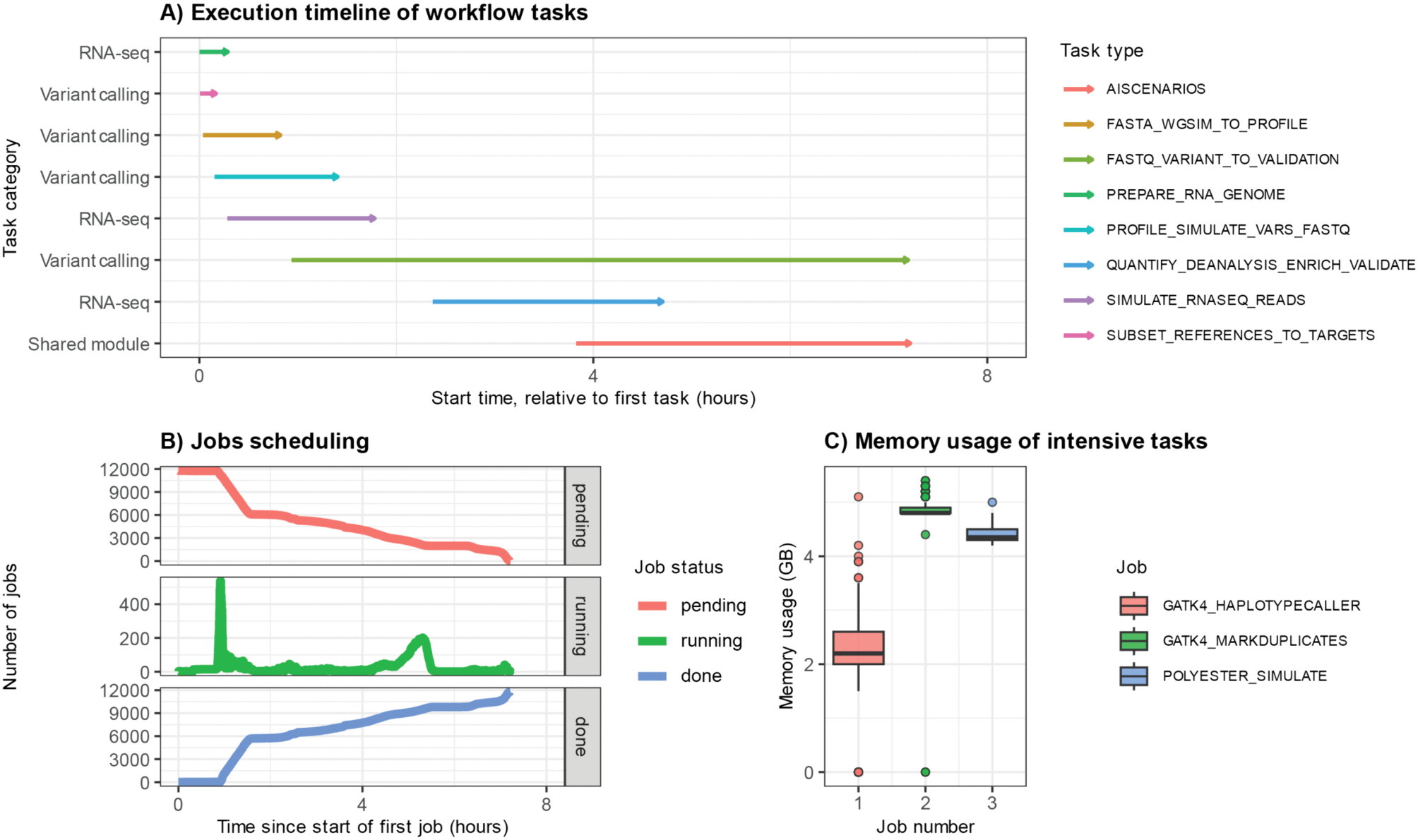
Progress and performance of the benchmark analysis on chromosome 22. (A) Gantt plot showing the execution timeline of eduomics tasks. Each bar represents the start and completion time of individual tasks. (B) The plot represents the progress of the analysis on Google Cloud, illustrating the dynamic status of tasks (pending, running, completed) over time. (C) Boxplot summarising memory utilisation for the top three most memory-intensive steps in the workflow.

The pipeline comprises a series of well-known memory-intensive tasks, such as GATK HaplotypeCaller for variant calling, GATK MarkDuplicates for identifying and marking duplicate reads, and polyester for RNA-seq read simulation. As shown in Fig 2C, these components exhibit distinct memory usage profiles reflecting their computational nature. HaplotypeCaller displayed the greatest variability, with memory consumption ranging from approximately 2 GB to 6 GB, consistent with the dynamic allocation required for the local de novo assembly performed during variant calling. MarkDuplicates maintained a stable peak around 5 GB, corresponding to the handling of large BAM files during duplicate identification. Similarly, polyester, required around 4,5 GB of memory. Overall, these results confirm that eduomics efficiently manages a large number of parallel jobs across multiple compute nodes, ensuring a stable performance and a smooth execution despite the complexity of the pipeline (Fig 2 and S1 Fig.).

The variant calling workflow achieved a validation rate of 38,2%, validating 241 out of 631 injected pathogenic variants. The RNA-seq workflow reached a validation rate of 78,5%, with 11 out of 14 differential expression patterns successfully recovered after quantification and differential expression analysis (S2 Fig). Fig 3 provides two illustrative examples of simulations generated from the benchmark on chromosome 22.

**Fig 3.**
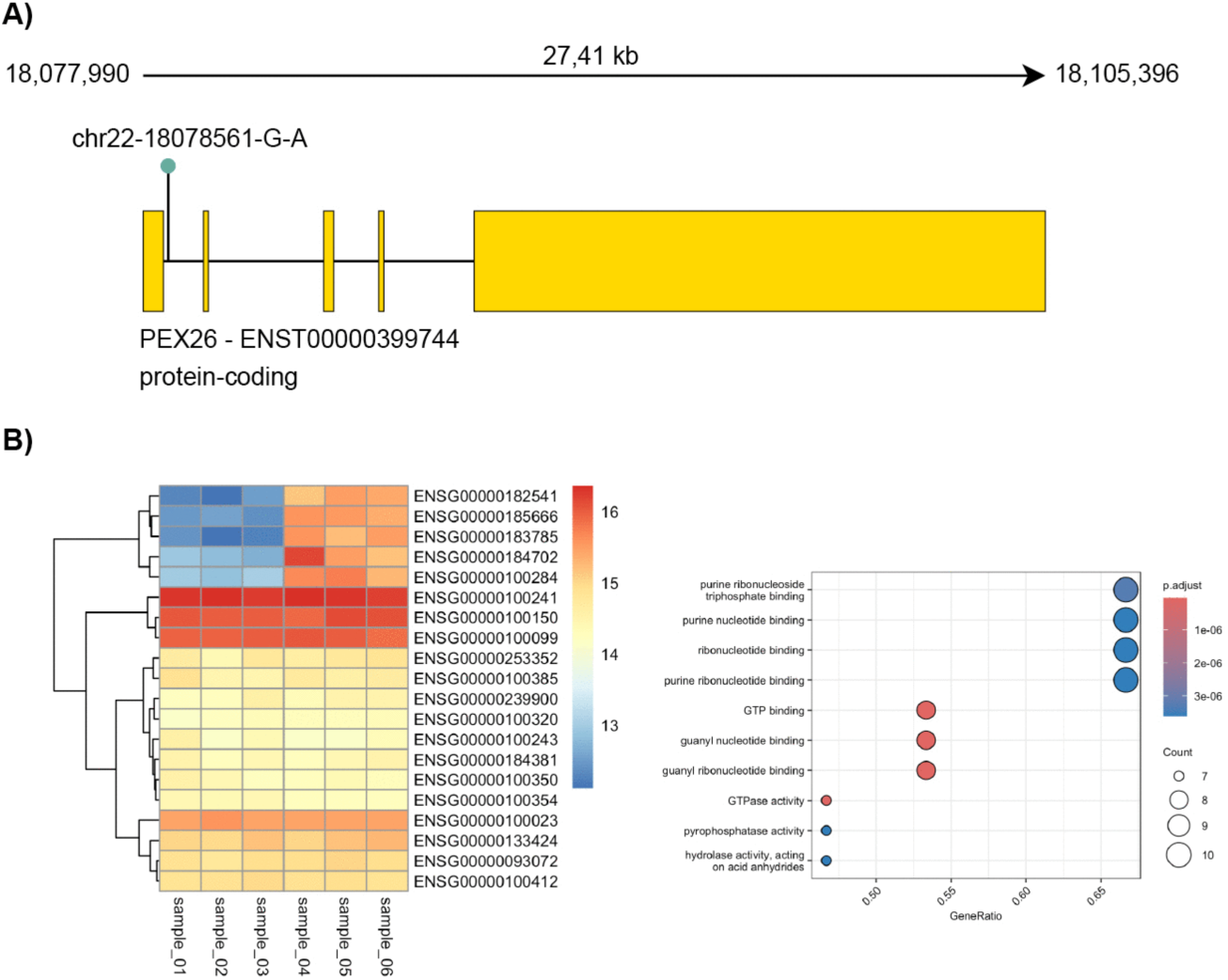
Exemplificative results from the benchmark analysis on chromosome 22. (A) Example from the DNA variant calling workflow. The sketch shows the *PEX26* gene (ENST00000399744) located on chromosome 22. In this simulation, a stop-gained mutation was introduced at position 18078561 (G>A). (B) Example from the RNA-seq workflow. Heatmap of the 20 most highly expressed genes across simulated samples, with colours ranging from blue (lower expression) to red (higher expression).

Fig 3A illustrates an example from the variant calling workflow. The case corresponds to the injected stop-gained mutation chr22-18078561-G-A in the *PEX26* gene (ENST00000399744), resulting in a premature termination codon (p.Trp62Ter). This variant is annotated as pathogenic/likely pathogenic in ClinVar. In the simulated dataset, aligned reads from the case sample display an adenine (A) at position 18078561, in contrast to the guanine (G) present in both the control sample and the human reference genome.

The affected codon lies within the N-terminal region of the peroxin-26 protein, a peroxisomal membrane anchor essential for the assembly and biogenesis of peroxisomes. Truncating mutations in *PEX26* are known to disrupt peroxisome biogenesis, leading to defective import of matrix proteins and the accumulation of very long-chain fatty acids (VLCFAs) [15]. The simulated clinical scenario (S1 Table) generated by eduomics reproduces a coherent genotype-phenotype correlation consistent with a peroxisome biogenesis disorder. The patient presents signs of developmental delay, hypotonia and early-onset pigmentosa. Consistent with the know literature on the disease [16,17], the generated clinical scenario describes biochemical investigations showing elevated VLCFA levels, which together with the neurological symptoms, visual impairment and liver involvement support the diagnosis of a peroxisomal biogenesis disorder. The integration of variant calling with the generation of a phenotypic and clinical history thus exemplifies the educational potential of the workflow in demonstrating the molecular link between a pathogenic variant and a real-world clinical scenario.

An example of the validated simulations generated by the RNA-seq workflow is presented in Fig 3B. The heatmap displays the 20 most highly expressed genes, highlighting a clear separation between control samples (sample 01-03) and case samples (sample 04-06). Notably, five genes (ENSG00000182541, ENSG00000185666, ENSG00000183785, ENSG00000184702, ENSG00000100284) shows a particular differential expression profile between conditions. These were subsequently identified as significantly differentially expressed, together with the complete set of significantly DEGs reported in S2 Table. The functional enrichment analysis revealed a strong over-representation of molecular functions related to nucleotide binding and GTPase activity (padj < 10⁻⁶), reflecting the involvement of genes in processes such as energy metabolism and intracellular signalling, which are frequently dysregulated in neurodevelopmental and skeletal disorders [18,19]. These findings support the biological coherence of the simulated dataset, which served as the molecular foundation for eduomics to generate an AI-derived clinical scenario. An example of such scenario is reported in S1 Table, illustrating a patient affected by progressive neurological decline, developmental delay, and skeletal abnormalities. The diagnostic investigation described in Gemini’s generated scenario revealed diffuse cerebral atrophy, ataxia and hypotonia, pointing to a complex neurological and skeletal disorder. In this AI clinical picture, whole-genome sequencing was also included among the diagnostic investigations to further elucidate the underlying molecular cause, mirroring real-world clinical practice. Similarly to the variant calling workflow, the integration of transcriptomic profiling with an AI-generated medical history exemplifies the educational potential of the RNA-seq workflow in linking altered gene expression patterns to clinically interpretable scenario.

## Discussion

Problem-based learning supported by data simulations provides students with the opportunity to practise data analysis in realistic and controlled environments. At the same time, such resources allow instructors to promote active learning through hands-on experience. Widespread introduction of simulated datasets in bioinformatics education is however hampered by several challenges. From a technical perspective, there are significant barriers to adoptions: steep learning curves, specific requirements each tool has, difficulties in deployment, lack of automation combined with the need to generate hundreds of dataset in contexts where simulated data serve also for tutoring and assessment [4–14]. From a pedagogical perspective, learning bioinformatics goes beyond acquiring methodological knowledge: however, we argue that problem-based learning alone is not sufficient to achieve long-term professional proficiency. According to the SOLO (structured of the observed learning outcome) taxonomy [20], higher-level intended learning outcomes (ILOs) involve the development of cognitive processes described by verbs such as “explain”, “theorise”, “hypothesise” [20,21]: achieving these outcomes requires students not only to move beyond the acquisition of procedural skills, but also beyond the application of specific tools to different questions and demonstrate instead a broader conceptual understanding. To reach these goals, data simulations must be integrated with a wider educational context that provides a clear purpose, an end-goal, an authentic motivation for the students to learn technical skills and apply them to real-life scenarios. At the same time, this context should train students to interpret their results in light of the bigger picture and to reach original conclusions. This pedagogical strategy creates a narrative framework that helps students understand not only how to perform analyses, but also the intrinsic behaviour of biological data they explore within a realistic context. We called this storyline-based learning.

Eduomics has the ambition to introduce a disruptive innovation in bioinformatics education by addressing these challenges and enabling such a holistic approach to teaching and learning. By developing this Nextflow-based pipeline, and automate all steps of NGS simulations, we demonstrate that realistic genomic and transcriptomic simulations can be made accessible to bioinformatics educators without prior knowledge of simulation software, and can be deployed in a scalable way, producing pedagogically meaningful datasets. With this solution, instructors are only required to define a chromosome and select the type of simulation: this approach eliminates technical entry barriers, and allows the adoption of a scenario-based teaching design with appropriate datasets. By generating hundreds of data combined with unique clinical scenarios, educators can assign unique datasets to each student or group, creating personalised learning experiences while maintaining reproducibility and consistency. This design supports a wide scale of teaching opportunities, from small-group workshops to large online courses. In addition, because each dataset is biologically validated against a known ground truth, instructors retain the full control over the simulated events and can ensure that analytical outcomes are always traceable back to their molecular causes. Thanks to its Nextflow implementation, eduomics is fully portable across infrastructures and can be deployed both on local high-performance clusters and in cloud environments at minimal cost. This ensures that high-quality simulations are affordable and accessible to educational institutions regardless of their computational resources.

Despite all these advantages, the current version of eduomics still presents a few limitations which will be addressed in future releases. The number of simulations produced by the variant-calling workflow is significantly higher compared to those generated by the RNA-seq workflow. This difference is partly expected, as the simulations were limited to a single chromosome. While each single variant can produce a simulation, genes must be selected in sets to simulate meaningful differential expression patterns. Therefore, the number of variants per chromosome is inherently larger that the number of genes sets which can be generated, and produce valid functional enrichments. The complexity of generating transcriptomic profiles is constrained by the gene ontologies of the selected chromosome. A key parameter influencing the outcome of the RNA-seq simulations is the similarity threshold (default = 0.3). By fine-tuning this parameter across different runs of the pipeline, users can modulate the number of the generated RNA-seq datasets.

In summary, eduomics bridges the gap between mastering computational tools and understanding their biological implications. It transforms complex simulation workflows into guided, narrative-driven learning experiences that allow educators to deliver, and students to appreciate, the richness and complexity of biomedical data. By combining automation with biological and clinical storytelling, eduomics redefines bioinformatics training into an engaging, creative and intellectually rewarding experience for the next generation of scientists.

## Materials and methods

### Pipeline implementation

Eduomics was implemented using the Nextflow DSL2 language [22], following the modular structure and best-practice guidelines of the nf-core community [23]. The workflow is composed of 16 modules from the shared nf-core repository and 14 modules specifically developed to enable the pipeline’s peculiar functionalities. These latter modules wrap custom scripts implemented in R, Bash and Python. Software dependencies are resolved using Conda, Docker, or Singularity, ensuring portability and reproducibility across computational environments. Continuous integration (CI) tests are executed through GitHub Actions on small datasets, while full-scale benchmarking was carried out on Google Cloud. The eduomics pipeline is openly available at https://github.com/lescai-teaching/eduomics.

### Input definition

The pipeline requires a sample sheet in CSV format, used to define the simulation parameters. This file specifies the type of workflow to be executed (variant calling or RNA-seq) and includes optional and mandatory fields. Among the optional parameters, two are particularly relevant: (i) capture, a path to an exome capture BED file, required for variant calling simulations; and (ii) the similarity threshold, a Jaccard index used to construct similarity networks from Gene Ontology annotations (see later), applied to RNA-seq simulations. Additional parameters allow users to select the chromosome, sequencing coverage, number of replicates, and grouping of simulated datasets.

### Variant calling simulation workflow

Reference sequences from the GRCh38 genome assembly [24] were subset to the user-specified chromosome and indexed using nf-core modules samtools_faidx (samtools v1.21) [25], bwa_index (bwa v0.7.18) [26], and gatk4_createsequencedictionary [27]. Target regions for the simulation were processed with the local module subsetcapture, implemented in Bash and relying on bedtools [28] and htslib [29]. This module compressed and indexed the capture BED file, extracted the chromosome-specific intervals, and generated padded regions (±50 bp and ±500 bp). Reference VCFs from gnomAD, Mills indels, 1000G, and dbSNP [24] were subset to the same regions using gatk4_selectvariants, providing interval-specific files. ClinVar variants have been downloaded from release GRCh38 in VCF format, and were processed with the local module subvar, implemented in Bash with htslib, which standardised chromosome nomenclature, extracted variants overlapping the target regions, and stratified the resulting set into distinct VCF files containing benign or pathogenic records. Together, these steps generated the DNA reference bundle, including a chromosome-specific genome FASTA with the relative index, the bwa index, capture targets, interval-specific VCFs, and curated ClinVar sets, which were used as the basis for read simulation and variant calling. Read simulation was carried out by first generating a profile for a sequencing experiment on the target chromosome with wgsim [30]: this generated paired-end FASTQ files from the chromosome-specific reference FASTA. The simulated reads were aligned to the same reference using bwa-mem, and the resulting BAM files were indexed with samtools. Downstream pre-processing followed the GATK Best Practices guidelines: duplicate reads were identified and marked with gatk4_markduplicates; base quality score recalibration (BQSR) was performed with gatk4_baserecalibrator, using known sites from dbSNP and Mills indels; the recalibrated BAMs were produced with gatk4_applybqsr and re-indexed; variants were called with gatk4_haplotypecaller. Finally, the resulting VCF was passed, together with the recalibrated BAM files and the chromosome-specific FASTA, to the local module simuscop_seqtoprofile. The module, implemented with SimuSCoP v1.1.2 [8], produced a profile file representing sequencing- and variant-related characteristics. The benign and pathogenic variant sets obtained from ClinVar were then converted into the variation format required by SimuSCoP using the local module pyconvertosim. This Python-based step (v3.9) generated three types of files: a baseline variation file containing only benign variants, a pathogenic variation file, and multiple combined variation sets, each including the benign background plus one pathogenic variant. Each combined variation file was then processed with the module simuscop_simureads, which integrates sequencing profiles, reference genome, capture intervals (± 500 bp), and the prepared variant sets to simulate case and control samples from the selected chromosome. For each pathogenic variant, paired-end FASTQ files were generated representing control samples (containing only benign variants) and case samples (containing both benign and the selected pathogenic variant).

Simulated case and control reads were then processed jointly following the GATK Best Practices, from alignment to variant calling producing single-sample gVCFs. To enable joint genotyping across samples, the gVCFs were processed with gatk4_genomicsdbimport and subsequently genotyped with gatk4_genotypegvcfs, generating a combined multi-sample VCF.

The combined VCF was then compared with the simulated case/control reads using the local module dnavalidation, implemented in Bash. This step verified the presence of the injected pathogenic variant within the simulated dataset by matching its genomic position in the VCF to the expected coordinates. The validated VCF together with the corresponding simulated reads were then stored in variant-specific result directories.

### RNA-seq simulation workflow

The GRCh38 genome assembly [24] was first subset and indexed to match the target chromosome with the nf-core module samtools_faidx. The genome GFF3 annotation from GENCODE v48 was filtered to retain only protein-coding transcripts from the user-selected chromosome. The annotation file was processed with the local module subsetgff, based on an R script relying on rtracklayer v1.66.0 [31] for GFF3 parsing, AnnotationDbi v1.68.0 [32] and org.Hs.eg.db v3.20.0 [33] to retrieve Gene Ontology (GO) annotations. To generate simulations in which differentially expressed genes produced a meaningful enrichment analysis, we selected a subset of transcripts that shared membership across enough ontologies and use these sets to generate the data. To this purpose, a transcript-GO incidence matrix was generated, a pairwise transcript similarity was calculated as a Jaccard index (default 0.3) [34], and a transcript similarity network was constructed with igraph v1.5.1 [35]. Gene communities were identified using the Louvain community detection algorithm, and to ensure biological plausibility, only clusters containing more than three but fewer than 10% of the total genes in the chromosome were retained. The biological relevance of these clusters was subsequently assessed through GO enrichment analysis performed with clusterProfiler v4.14.0 [36], producing a set of gene lists. The reference transcriptome (GENCODE v48 FASTA) was filtered with the local module subsetfastatx, which implements an R script for transcript-annotation matching and uses Biostring v2.74.0 [32] for sequence handling, resulting in a chromosome-specific transcript FASTA.

Synthetic count matrices were then produced with the local module countmatrices, based on in-house R functions. Baseline counts were initialised proportionally to transcript length and user-defined coverage, then fold changes were applied to the GO-selected transcripts to encode coherent up- and down-regulation across groups and replicates. Read-level simulation was carried out with the local module polyester_simulate, which uses the polyester package v1.38.0 [10] to generate synthetic reads from the filtered transcript FASTA. In the current implementation, the module operates in count-matrix mode, receiving the simulated count matrices as input to produce the reads. Although the workflow supports an alternative mode based on simulated fold changes, this functionality is not yet implemented. Simulated reads were produced in gzipped FASTA format and provided as input for downstream analyses. Quantification of simulated reads was performed with the nf-core module salmon_quant using salmon v1.10.3 [37] in mapping-based mode. Gene-level counts were then obtained in R with tximport v1.34.0 [38] by mapping transcripts to genes using the tx2gene file derived from the filtered annotation. Differential expression analysis was carried out through the deanalysis module, implemented using DESeq2 v1.46.0 [39]. The sample design was constructed from the specified numbers of replicates and groups, genes with low counts were pre-filtered, and the condition was modelled using the design formula ∼ condition. Standard diagnostic (including MA, dispersion, and counts plot) and exploratory summary plots (PCA and heatmap) were generated. The results were exported as a ranked table listing adjusted P-values (padj) and log₂ fold changes. Genes were defined as differentially expressed when padj < 0.05. Functional enrichment was performed using the enrichment module with clusterProfiler, testing the different GO ontologies (BP, MF and CC). Summary visualisations (dot plots and cnetplots) were generated for each ontology. A final validation step was implemented in the local module rnaseqvalidation which relies on an R script using DOSE v4.0.0 [40]. A simulation was classified as successful when at least one ontology (BP, MF, or CC) contained three or more significantly enriched terms. Upon validation, the RNA framework produced a complete set of outputs suitable for downstream training and interpretation. These include: (i) a reference bundle (FASTA transcriptome, index files, annotation GFF3), (ii) simulated reads in gzipped FASTA format, (iii) differential expression results, (iv) functional enrichment results and (v) a collection of diagnostic and exploratory plots.

### AI-based scenario generation

A final module, aiscenarios, is shared by both the variant calling and RNA-seq workflows. This module generates patient-inspired clinical narratives from the simulated data using the Google Gemini API. Depending on the input, the module ingests either pathogenic variants with the corresponding gene from the variant calling workflow or lists of differentially expressed genes from the RNA-seq workflow. The corresponding Python script, implemented with the google-genai library (v1.9.0) [41], instructed Gemini to generate a detailed clinical scenario describing a hypothetical patient, including relevant clinical features, family history, and diagnostic clues consistent with the underlying molecular and genetic dysfunctions. The output of this module consisted of a text file containing realistic clinical cases, which serve as the final layer of the storyline-based educational framework of eduomics.

## Acknowledgments

We acknowledge Harshil Patel, Phil Ewels, Friederike Hanssen, Gavin Mackenzie, Jeremy Guntoro, Kevin Menden, Matthias De Smet, Maxime U Garcia, Michael J Cipriano, Patrick Hüther, Priyanka Surana, Ramprasad Neethiraj, Santiago Revale, Suzanne Jin and the nf-core community for developing community modules used in eduomics.

## Authors’ Contributions

Conceptualization: FL; Methodology: FL, LS, DB, SC; Software: LS, FL, DB, SC, MS; Formal Analysis: LS; Writing original draft: LS, FL; Writing review and editing: LS, FL, DB; Visualization: LS, SC.

## Supporting information

**S1 Fig. Metromap of the eduomics pipeline.** The eduomics pipeline implements two different workflows, variant calling (in blue) and RNA-seq (in amaranth). Both rely on a configuration CSV file that defines the input data and the type of simulation to be performed. Each path follows a series of sequential steps that transform raw data into validated results according to defined quality criteria. The results of both workflows converge in a shared module implemented with Google Gemini API, which generates realistic patient-inspired clinical scenarios based on the simulated variants or the lists of differentially expressed genes.

**S2 Fig. Validation rate of the eduomics pipeline on chromosome 22.** Infographic summarising the validation outcomes of eduomics simulations in the benchmarking study on chromosome 22.

**S1 Table. Example of simulated clinical scenarios for variant calling and RNA-seq workflows.** The table presents two informative, patient-inspired clinical scenarios generated through the integration of the Google Gemini API within eduomics. Each scenario includes detailed information such as patient presentation, symptoms, family history, and a possible diagnosis.

**S2 Table. Differentially expressed genes in the use-case RNA-seq simulation.** List of differentially expressed genes identified in the illustrative RNA-seq simulation. Each entry includes the corresponding Ensembl Gene ID and Gene Symbol.

## Data availability

The code used in this study is publicly available on our GitHub repository https://github.com/lescai-teaching/eduomics. A DOI has been assigned to the repository via Zenodo: <u>10.5281/zenodo.15835069</u>.

