## Supplementary figures and images for "Eduomics: a Nextflow pipeline to simulate -omics data for education"

### Supplementary Figure 1

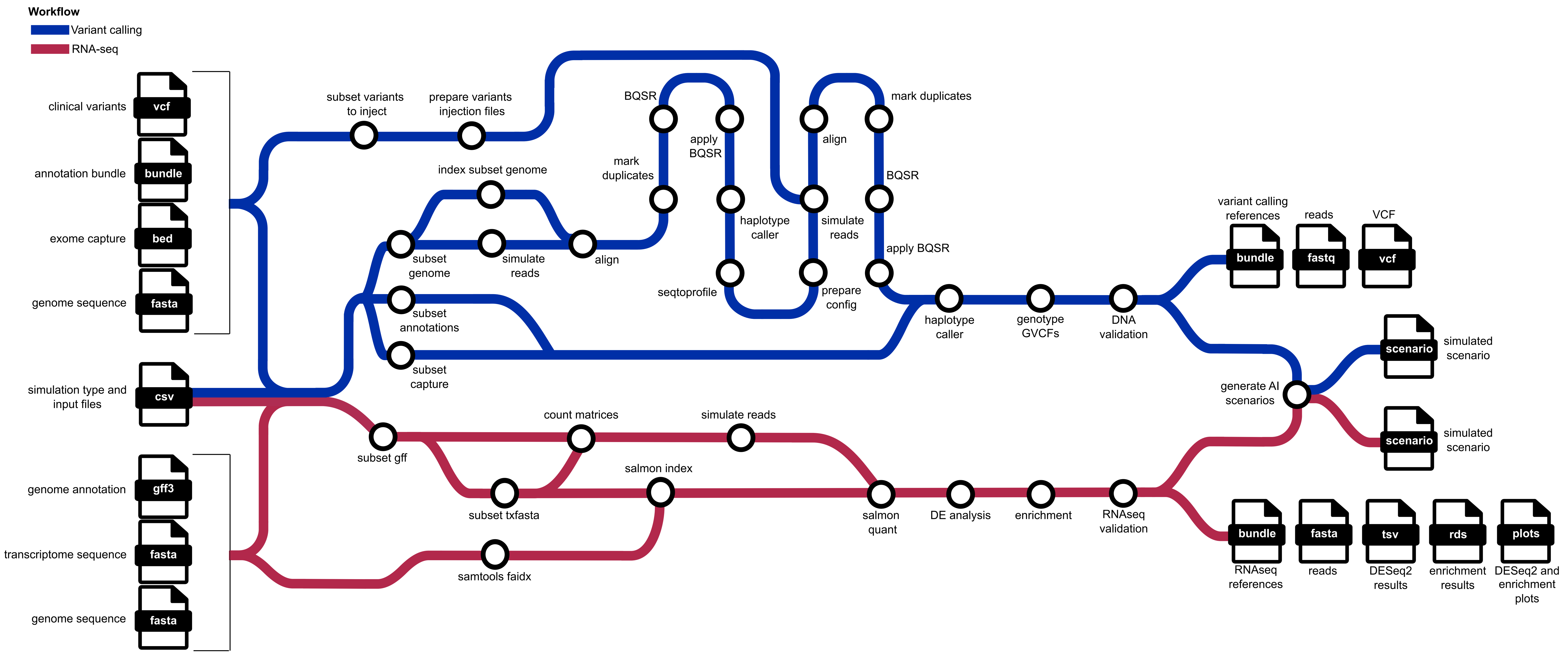

### Supplementary Figure 2

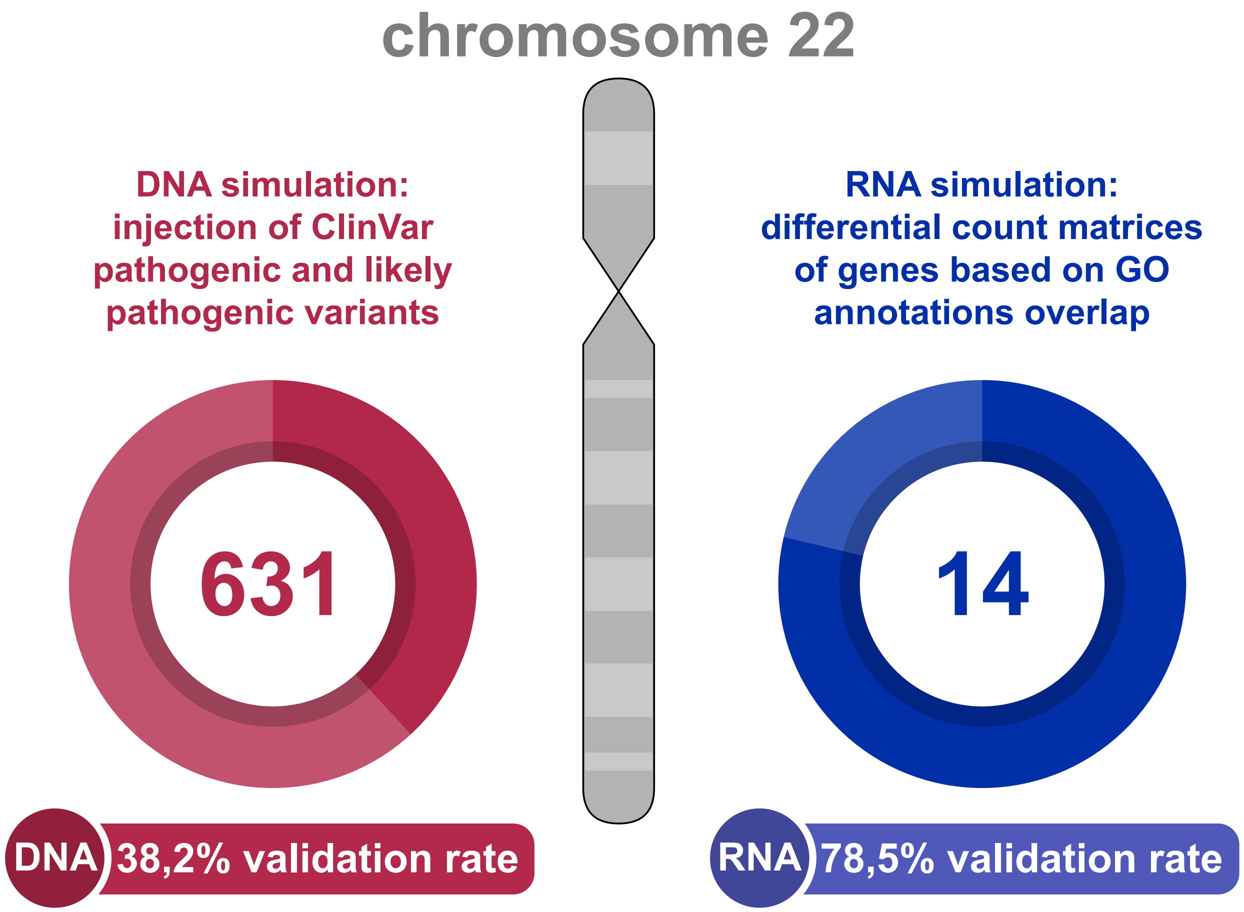
